# A novel Mendelian randomization method identifies causal relationships between gene expression and low-density lipoprotein cholesterol levels

**DOI:** 10.1101/671537

**Authors:** Adriaan van der Graaf, Annique Claringbould, Antoine Rimbert, BIOS consortium, Harm-Jan Westra, Yang Li, Cisca Wijmenga, Serena Sanna

**Affiliations:** University of Groningen, University Medical Centre Groningen, Department of Genetics, Groningen, the Netherlands; University of Groningen, University Medical Centre Groningen, Department of Pediatrics, Section Molecular Genetics, Groningen, the Netherlands; l’institut du thorax, Unité Inserm UMR 1087 / CNRS UMR 6291, IRS-UN, 8 Quai Moncousu, Nantes cedex1, France; Helmholtz Centre for Infection Research, Inhoffenstr. 7, 38124 Braunschweig, Germany; Centre for Individualised Infection Medicine, Feodor-Lynen-Str. 7, 30625 Hannover, Germany

**Author notes:** These authors jointly supervised this work (Yang Li, Cisca Wijmenga, Serena Sanna).

## Abstract

Robust inference of causal relationships between gene expression and complex traits using Mendelian Randomization (MR) approaches is confounded by pleiotropy and linkage disequilibrium (LD) between gene expression quantitative loci (eQTLs). Here we propose a new MR method, MR-link, that accounts for unobserved pleiotropy and LD by leveraging information from individual-level data. In simulations, MR-link shows false positive rates close to expectation (median 0.05) and high power (up to 0.89), outperforming all other MR methods we tested, even when only one eQTL variant is present. Application of MR-link to low-density lipoprotein cholesterol (LDL-C) measurements in 12,449 individuals and eQTLs summary statistics from whole blood and liver identified 19 genes causally linked to LDL-C. These include the previously functionally validated *SORT1* gene, and the *PVRL2* gene, located in the *APOE* locus, for which a causal role in liver was yet unknown. Our results showcase the strength of MR-link for transcriptome-wide causal inferences.

## Introduction

Mendelian randomization (MR) is a method that can infer causal relationships between two heritable complex traits from observational studies^1,2^. In recent years, MR has gained popularity in the epidemiological field and has provided valuable new insights into the risk factors that cause diseases and complex traits^1–3^. MR has, for example, successfully identified causal relationships between low-density lipoprotein cholesterol (LDL-C) and coronary artery disease, results which have informed therapeutic strategies^4,5^. MR analysis has also shown that a causal relationship between high density lipoprotein cholesterol (HDL-C) and coronary artery disease is unlikely, which is in contrast to previous epidemiological associations^6^. The same approach has been applied to identify molecular marks that are causal to disease^7–10^. Since gene expression is one of these marks, investigating its causal role in complex traits is of particular interest given that complex trait loci are enriched for expression quantitative trait loci (eQTLs)^11^.

MR relies on certain assumptions to correctly infer a causal relationship between an exposure (e.g. a risk factor) and an outcome (e.g. a complex trait). MR can leverage QTL variants of the exposure as instrumental variables (IVs) if three conditions are met: the IVs have to be i) associated with the exposure, ii) independent of any confounder of the exposure-outcome association and iii) conditionally independent of the outcome given the exposure and confounders. One major challenge of applying MR to gene expression is correcting for deviations from the third assumption, which can occur in the presence of linkage disequilibrium (LD) between the eQTLs used as IVs, or in the presence of pleiotropy, i.e. when IVs affect the outcome through pathways other than the exposure of interest. Accounting for LD is necessary when gene expression is the exposure trait in MR because, in contrast to the majority of complex traits, the genetic architecture of gene expression is characterized by the presence of strong-acting eQTLs located proximal to their transcript (i.e. in *cis*), which are often correlated through LD^12,13^. Similarly, the presence of pleiotropy cannot be excluded *a priori* given that the majority of variants in our genome are likely to affect one or multiple phenotypes^14–16^. While there are MR-methods^7,8,10,17–21^ that extend standard MR analysis to correct for LD and pleiotropy, the application of these methods is not optimal because they require either the removal of pleiotropic IVs from the statistical model^7,20,21^ or that all sources of pleiotropy are measured and incorporated into the model^22,23^. These constraints limit robust inference of the causal role of gene expression traits as there is often only a limited number of IVs (i.e. eQTLs) available, and subsequent removal of outliers will substantially reduce power. Likewise, it is not always possible to measure all sources of pleiotropy because pleiotropy could come from expression of a gene in a different tissue or even from other unmeasured molecular marks or phenotypes.

Here we introduce MR-link, a novel two-sample MR method that allows robust causal inference in the presence of LD and unobserved pleiotropy, and without requiring the removal of pleiotropic IVs or measuring all sources of pleiotropy. MR-link uses summary statistics of an exposure combined with individual-level data on the outcome to estimates the causal effect of an exposure from IVs (i.e. eQTLs if the exposure is gene expression) while at the same time correcting for pleiotropic effects using genetic variants that are in LD with these IVs (**Figure 1**).

**Figure 1.**
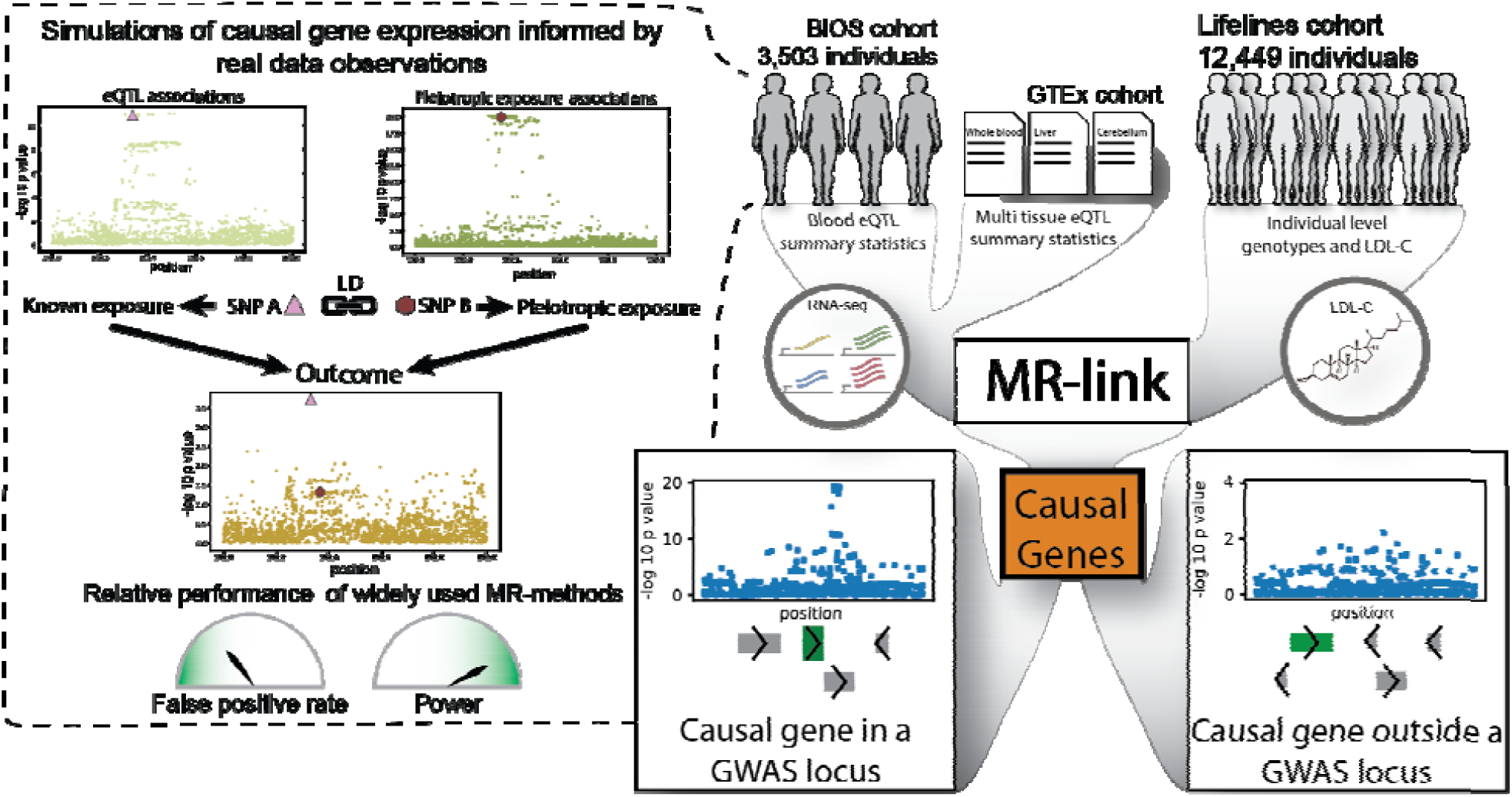
Graphical representation of the study. The Biobank Integrative Omics Study (BIOS) cohort was used to identify expression quantitative trait loci (eQTL) and characterize the genetic architecture of gene expression. Dashed outbox: Knowledge used in a simulation scheme that mimicked gene expression traits, including linkage disequilibrium (LD) between eQTL. We used this simulation to assess the false positive rates (FPR) and power for widely used Mendelian randomization (MR) methods. We applied our new MR-method, MR-link, to both the simulations and to individual-level data of low density lipoprotein cholesterol (LDL-C) in 12,449 individuals (Lifelines) and used BIOS and GTEx eQTL summary statistics to identify gene-expression changes causal to LDL-C within or outside a genome-wide association study (GWAS) locus.

We assessed the performance of MR-link using simulated data in 100 different scenarios that mimicked the genetic architecture of gene expression, information we derived by looking at eQTL association patterns in a large cohort of samples with genetic and transcriptomics data^13^. We then applied MR-link to individual-level data for LDL-C measurements in 12,449 individuals with four different eQTL summary statistic data sets: blood eQTLs identified in the BIOS cohort (**Figure 1**) and eQTLs from blood, liver and cerebellum identified by the GTEx Consortium^24^ (**Figure 1**). Our results in simulated and real data show that MR-link can robustly identify causal relationships between gene expression and an outcome (e.g. a complex trait), even when the information for causal inference is very limited (i.e. only a single instrumental variable is available).

## Results

### Genetic regulation of gene expression is often shared between genes through linkage disequilibrium

To investigate how the genetic effects on gene expression are distributed in *cis*, we searched for eQTLs 1.5 megabases (Mb) on both sides of the translated region of 19,960 genes (**Methods**). We used data from the BIOS cohort, a cohort of 3,503 Dutch individuals whose genome and whole blood transcriptome has been characterized (**Figure 1**), and applied a summary statistics–based stepwise linear regression approach (GCTA-COJO) to identify jointly significant variants, e.g. one or more variants that jointly associate significantly with expression changes of a gene^13,25^ (**Methods**). We observed that 58% of the genes with an eQTL at *p* < 5×10^−8^ (10,831 genes) had at least one jointly significant eQTL at *p* < 5×10^−8^ (**Methods**) (**Figure 1** and **Figure 2A**). These genetic effects were mostly non-overlapping: only 7.0% of the genes have the exact same top eQTL variant. In contrast, genetic variants regulating gene expression of a gene were very often in LD with other eQTL: 30% of top variants are in LD (r^2^ > 0.5) between genes, and this percentage increased to 58% if all jointly significant eQTLs are considered (**Methods**). These observations suggest that presence of pleiotropy between eQTLs used as IV in transcriptome-wide MR analyses are mostly attributable to variants in LD with the IVs (pleiotropy through LD), rather than to completely overlapping IVs (pleiotropy through overlap) (**Figure 2**).

**Figure 2.**
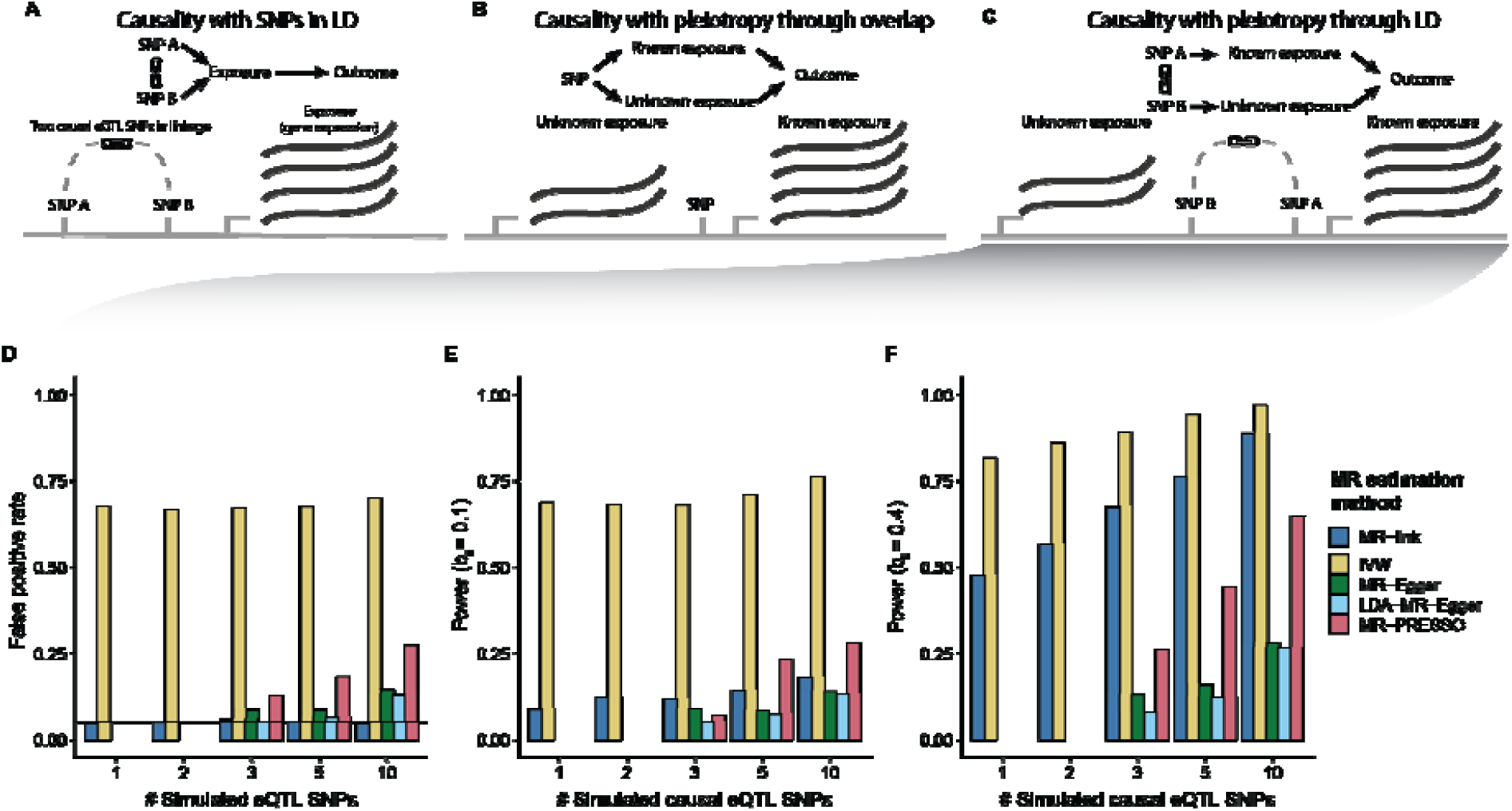
Typical scenarios of pleiotropy in causal inference using gene expression and relative performance of different MR methods. The top three panels display typical scenarios to consider when performing causal inference in gene expression: (**A**) instrumental variables (IVs) for the same gene (exposure) are in linkage disequilibrium (LD), (**B**) pleiotropy is present through overlap of IVs (pleiotropy through overlap) and (**C**) pleiotropy is present through LD between IVs for different exposures (pleiotropy through LD). Bottom panels show simulation results under the pleiotropy through linkage scenario when 1, 3, 5 or 10 causal SNPs were simulated. (**D**) False positive rates (at alpha = 0.05) in scenarios where no causal relationship is simulated. (**E**) Power to detect a small causal effect (at alpha = 0.05). (**F**) Power to detect a large causal effect (at alpha = 0.05). Note that MR-link is the only MR method that can adjust for pleiotropy when only one or two instruments are available. MR methods that had fewer than 100 out of 1,500 estimates in a scenario are not shown (**Methods**). Extended results depicted in panels **D-F** can be found in **Supplemental Table 2**.

### MR-link outperforms other MR methods in false positive rate and power

Based on the observation that genetic variation of gene expression is mostly characterized by eQTLs that are in LD, but not overlapping between genes, we developed a novel MR method that infers causality while accounting for pleiotropy and LD (**Methods, Figure 2, Supplemental Note and Supplementary Figure 1**). We assessed the performance of MR-link and compared it to four other MR methods: Inverse variance weighting (IVW), which assumes absence of LD and pleiotropy, and the pleiotropy-robust methods MR-Egger, LDA-MR-Egger and MR-PRESSO (**Table 1**)^17,18,20,26^. Here we simulated causal relationships between an exposure and an outcome in a 5Mb region, based on LD structure estimated for 403 European samples from the 1000 Genomes project^27^ (**Methods**). We assessed all tested MR-methods in 1,500 simulated data sets for 100 different scenarios, reflecting absence of causality, moderate to high causal effects, coupled with presence or absence of pleiotropic effects and different number of IVs. We initially evaluated two approaches to select QTL variants as IVs: GCTA-COJO and *p* value clumping (**Methods**)^25,28^. We observed that GCTA-COJO was best suited for IV selection because: (i) the median number of IVs identified by GCTA-COJO better represented the number of simulated causal variants (**Supplemental Table 1**) and (ii) the false positive rates (FPR) in the MR analysis using the IVW method was lower (median FPR was 0.057 using GCTA-COJO versus 0.115 using clumping) (**Supplemental Figure 1** and **Supplemental Table 1**). We therefore selected IVs for the exposure using the GTCA-COJO approach in subsequent analyses.

**Table 1.** MR Methods assessed in this study. MR methods assessed in our simulation, with information about the type of data needed to make a causal estimate, ability to correct for pleiotropy or linkage disequilibrium and the minimum number of instrumental variables required. Abbreviations: IVW: inverse variance weighting, sumstats: summary statistics, ILD: individuals level data, O: outcome, E: Exposure, LDR: linkage disequilibrium reference (LDR) panel. For more details, see **Methods**. ^a^ We refer here to the MR-PRESSO test that reports estimates after identifying and removing outliers, as the test without outliers generalizes to IVW estimates.

| MR method | Data required | Pleiotropy correction | LD correction | Minimum no. of IVs required |
| --- | --- | --- | --- | --- |
| MR-link | Sumstats: E, ILD: O | Yes | Yes | 1 |
| Inverse variance weighted (IVW)<sup>26</sup> | Sumstats: O, E | No | No | 1 |
| LDA-MR-Egger<sup>17</sup> | Sumstats: O, E, LDR | Yes | Yes | 3 |
| MR-Egger<sup>18</sup> | Sumstats: O, E | Yes | No | 3 |
| MR-PRESSO<sup>a,20</sup> | Sumstats: O, E | Yes | No | 3<sup>a</sup> |

**Table 2.** The 15 genes identified by MR-link as causal for LDL-C levels in the analysis that included eQTLs from the BIOS cohort. Gene names are according to ENSEMBL GENES 96 database (human build 37). The causal effect estimate represents the changes in LDL-C (mg/dL) per standard deviation increase in gene expression. Full summary statistics of the genes are shown in **Supplemental table 4**. Abbreviations: LDL-C: Low density lipoprotein cholesterol, HDL-C: High-density lipoprotein cholesterol GWAS: Genome wide association study, siRNA: small interfering RNA

| Gene Name | Estimated causal effect | p value | #IVs | Biological function and link to LDL-C |
| --- | --- | --- | --- | --- |
| *IGLC5* | -0.0531 | 7.85E-10 | 1 | Immunoglobulin lambda constant 5 (*IGLC5*) is a pseudogene; two other genes in the same locus and belonging to the same family appear in this table (*IGLV4-69* and *IGLC6*). *IGLC5* does not have a known function in cholesterol metabolism |
| *KB-1460A1.5* | 0.1535 | 5.61E-09 | 1 | *KB-1460A1.5* is an RNA gene with unknown function in cholesterol metabolism |
| *MIR4482-1* | -0.1008 | 1.47E-08 | 3 | MicroRNA 4482 (*MIR4482-1*) is an RNA gene that does not have known function in cholesterol metabolism |
| *UNC5B* | -0.0124 | 2.15E-08 | 1 | Unc-5 Netrin Receptor B (*UNC5B*) is a protein-coding gene and a receptor for the *NETRIN1* protein. It is associated to the disease Hyperinsulinemic Hypoglycemia familial 3. *UNC5B* is localized in lipid rafts, membrane compartments that contain high levels of cholesterol and lipids<sup>50</sup>. Higher *UNC5B* expression levels are associated to insulin resistance<sup>36,50,51</sup>. While insulin resistance can be related to hypercholesterolemia, the direct link of this gene to LDL-C remains unclear. |
| *TMEM176B* | -0.0287 | 5.16E-08 | 4 | Transmembrane protein 176B (*TMEM176B*) is a protein coding gene located in the *TMEM176B - TMEM176A-AOC1* GWAS locus for HDL-C<sup>31</sup>. Two IVs for *TMEM176B* are overlapping with IVs for *TMEM176A*. |
| *REEP1* | -0.0218 | 5.78E-08 | 1 | Receptor Accessory Protein 1 (*REEP1*) is a protein-coding gene. Mutations of the N-terminal of *REEP1* lead to accumulation in lipid droplets in the endoplasmic reticulum<sup>52</sup>. *REEP1* does not have a known function in cholesterol metabolism. |
| *IGLC6* | -0.0768 | 1.18E-07 | 3 | Immunoglobulin lambda constant 6 (*IGLC6*) is a pseudogene; two other genes in the same locus belonging to the same family appear in this table (*IGLV4-69* and *IGLC5*). *IGLC6* does not have a known function in cholesterol metabolism. |
| *AOC1* | -0.0086 | 1.19E-07 | 1 | Amine Oxidase, Copper Containing 1 (*AOC1*) is a protein-coding gene located in the *TMEM176A-TMEM176B-AOC1* GWAS locus for HDL-C<sup>31</sup>. |
| *IGLV4-69* | 0.0877 | 1.51E-07 | 1 | Immunoglobulin Lambda Variable 4-69 (*IGLV4-69*) is a protein-coding gene; two other genes in the same locus and belonging to the same family appear in this table (*IGLC5* and *IGLC6*). *IGLV4-69* does not have a known function in cholesterol metabolism |
| *SYCP2L* | -0.0194 | 2.02E-07 | 2 | Synaptonemal Complex Protein 2 Like (*SYCP2L*) is a protein-coding gene, important in oocyte cell survival, but also highly expressed in neutrophils. *SYCP2L* is located in a GWAS locus for antiphospholipid syndrome and fatty acid measurements. Functionally, not much is known about *SYCP2L*, and it does not have a known function in cholesterol metabolism<sup>35,53</sup>. |
| *C10orf10/DEPP1* | -0.0689 | 2.35E-07 | 1 | *DEPP1* (also known as *C10orf10*) is an autophagy regulator highly expressed in adipose tissue. *DEPP1* overexpression in mice reduces glucose and triglyceride levels<sup>54</sup>, although a direct link of this gene to LDL-C remains unclear. |
| *TMEM176A* | -0.0224 | 2.39E-07 | 3 | Transmembrane protein 176A (*TMEM176A*) is a protein-coding gene located in the *TMEM176B - TMEM176A-AOC1* GWAS locus for HDL-C<sup>31</sup>. Two IVs for *TMEM176A* are overlapping with *TMEM176B*. |
| ***RP11-18H21.1*** | 0.0296 | 2.50E-07 | 2 | *RP11-18H21.1* is a non-coding RNA gene without a known function in cholesterol metabolism. |
| ***TACSTD2*** | -0.0186 | 4.48E-07 | 2 | Tumor Associated Calcium Signal Transducer 2 (*TACSTD2*) is a protein-coding gene located on the cell membrane involved in the superpathway "Ca, cAMP and Lipid Signaling". The function of *TACSTD2* in LDL-C metabolism is unclear. |
| ***PRDM5*** | 0.0666 | 2.73E-06 | 4 | PR/SET Domain 5 (*PRDM5*) is a protein-coding gene and a member of the PR family of transcription factors. In human colon cancer cells and *Prdm5* mutant mouse intestines, dysregulation of *PRDM5* affects monoacylglycerol lipase (*MGLL*) expression and other genes involved in lipid metabolism<sup>55</sup>. The direct link to LDL-C remains unclear. |

When we simulated pleiotropy through LD with no causal effect of the known exposure (gene) on the outcome (**Methods, Supplemental Table 2** and **Figure 2C, 2D**), all existing MR-methods showed inflated FPR (up to 0.71, 0.15, 0.13 and 0.27 for IVW, MR-Egger, LDA-MR-Egger and MR-PRESSO, respectively), whereas MR-link presented an FPR close to expectation (median: 0.05, maximum: 0.058). In addition, for LDA-MR-Egger, MR-Egger and MR-PRESSO, the FPR was undesirably dependent on the number of causal SNPs simulated (**Figure 2D**).

In the scenarios of pleiotropy through LD and non-null causal effects (*b*_*E*_ =0.05, *b*_*E*_ =0.1, *b*_*E*_ =0.2 and *b*_*E*_ =0.4), MR-link has high detection power (up to 0.89) and strongly outperforms all other pleiotropy-robust methods (maximum detected power was 0.28 for MR-Egger, 0.26 for LDA-MR-Egger and 0.65 for MR-PRESSO) (**Figure 2E-F, Supplemental Table 2** and **Methods**). Among all the methods tested, including MR-link, and for all scenarios, IVW had the greatest detection power but also an inflated FPR (minimum FPR: 0.63), making this MR method unsuitable in such pleiotropic scenarios (**Methods**).

When we simulated pleiotropy through LD for a large subset of the simulated causal SNPs (**Figure 2B** and **Methods**), a situation we expect to be rare in real-world scenarios based on our observation in the BIOS cohort, we observed that all methods including MR-link have increased FPR (up to 0.22 for MR-link, 0.77 for IVW, 0.10 for LDA-MR-Egger, 0.13 for MR-Egger and 0.30 for MR-PRESSO) (**Supplemental Table 3**). Nonetheless, MR-link remains a powerful method when a causal effect is simulated: maximum power was 0.79 for MR-link, 0.98 for IVW, 0.29 for MR-Egger, 0.28 for LDA-MR-Egger and 0.65 for MR-PRESSO (**Supplemental Table 3**). Although IVW here resulted, again, in the highest power (0.98), the FPR was likewise largely inflated (0.77)

**Table 3.** The four genes identified by MR-link as causal for LDL-C levels in the analysis that included eQTLs from liver tissue in the GTEx study. Gene names according to ENSEMBL GENES 96 database (human build 37). The causal effect estimate represents changes in LDL-C (mg/dL) per standard deviation increase in gene expression. Full summary statistics of these genes are shown in (**Supplemental table 5**). Abbreviations: LDL-C: Low density lipoprotein cholesterol, GWAS: Genome wide association study, siRNA: small interfering RNA

| Gene Name | Estimated causal effect | p value | #IVs | Biological function and link to LDL-C |
| --- | --- | --- | --- | --- |
| **PVRL2** | 0.3177 | 3.0E-14 | 1 | Poliovirus Receptor-Related 2 (PVRL2) is a protein-coding gene also known as *NECTIN2*. It is a cell-membrane protein located in the LDL-C GWAS locus *APOE*. siRNA experiments show that LDL-C uptake is increased in cells upon its downregulation<sup>44,56</sup>. PVRL2 knockout mice also had less atherosclerosis<sup>42</sup>. Both studies indicate a reduction in LDL-C upon downregulation of *PVRL2*. |
| **PSRC1** | -0.0847 | 3.99E-09 | 1 | Proline And Serine Rich Coiled-Coil 1 (*PSRC1*) is a protein-coding gene located in an LDL-C GWAS locus<sup>31</sup>. *PSRC1* has not been found to have an effect on cholesterol despite being targeted in a specific functional study<sup>40</sup>. The IV for *PSRC1* is overlapping with the IV for *CELSR2* and *SORT1*. |
| **SORT1** | -0.0865 | 5.89E-09 | 1 | Sortilin (*SORT1*) is a protein-coding gene located in an LDL-C GWAS locus<sup>31</sup>. siRNA and knockdown experiments have functionally validated that *SORT1* has a negative effect on LDL-C levels<sup>38,39,57</sup>. The IV for *SORT1* is overlapping with the IV for *PSRC1* and *CELSR2*. |
| **CELSR2** | -0.0993 | 6.8E-08 | 1 | Cadherin EGF LAG Seven-Pass G-Type Receptor 2 (*CELSR2*) is a protein-coding gene located in an LDL-C GWAS locus<sup>31</sup>. *CELSR2* has not been found to have an effect on cholesterol despite being targeted by a specific functional study<sup>40</sup>. The IV for *CELSR2* is overlapping with the IV for *PSRC1* and *SORT1*. |

### MR-link identifies known and novel causal relationships between gene expression changes and LDL-cholesterol levels

We applied MR-link to four separate summary statistics–based eQTL datasets combined with individual-level genotype data and LDL-C measurements in 12,449 individuals from the Lifelines cohort^29^ (**Figure 1**). We assessed the causal effect of gene expression changes in i) whole blood using eQTLs from BIOS (n=3,503) and GTEx (n=369), ii) liver as the main tissue important for cholesterol metabolism (using eQTLs from GTEx, n=153), and iii) cerebellum tissue (using eQTLs from GTEx,n=154) as a tissue not involved in cholesterol metabolism but with similar sample size (and thus power) to liver tissue^24,30^.

Transcriptome-wide application of MR-link to these eQTL datasets identified 21 significant genes whose variation in blood (15 using BIOS eQTLs, 2 using GTEx eQTLs) or liver (4 genes) was causally related to LDL-C (**Table 2, Table 3, Supplemental Table 4** and **Supplemental Table 5**). No significant genes were found in cerebellum (**Supplemental Table 5**).

MR analysis that used whole-blood eQTLs from GTEX was, as expected, underpowered compared to the analysis using BIOS eQTLs. Only two genes were found to be significant here, but they were non-significant in the BIOS cohort, where a more robust estimate could be made thanks to higher number of IVs identified (**Supplemental Figure 2A**). Despite the limited power, we observed high concordance between effect sizes from the two analyses (using blood eQTLs from BIOS and GTEx) for all genes that showed nominal significance (*p* < 0.05) in the BIOS analysis, with 89.3% of genes showing the same effect direction (**Supplemental Figure 2B**).

Several genes located in GWAS loci for cholesterol metabolism were found in the MR analysis that used blood eQTLs from BIOS. These include *AOC1, TMEM176A* and *TMEM176B*, which are all located in the same HDL-C-associated locus^31–33^, and *SYCP2L*, which is located in a GWAS locus for polyunsaturated fatty acids and related to LDL-C levels^34,35^. For the other genes identified, there was no direct evidence in the literature for a direct role in cholesterol metabolism, although some interesting patterns were evident. For example, we observed multiple genes involved in immunoglobulin production (*IGLC5, IGLC6* and *IGLV4-69*) and insulin metabolism (*UNC5B, DEPP1*), observations at least consistent with the role of cholesterol in inflammation and insulin resistance^36,37^. Of note, for all 15 genes, the effect direction estimated by MR-link was concordant with the direction estimated by other MR-methods (**Table 1, Table 2, Supplemental Figure 3** and **Supplemental Table 6**).

**Figure 3.**
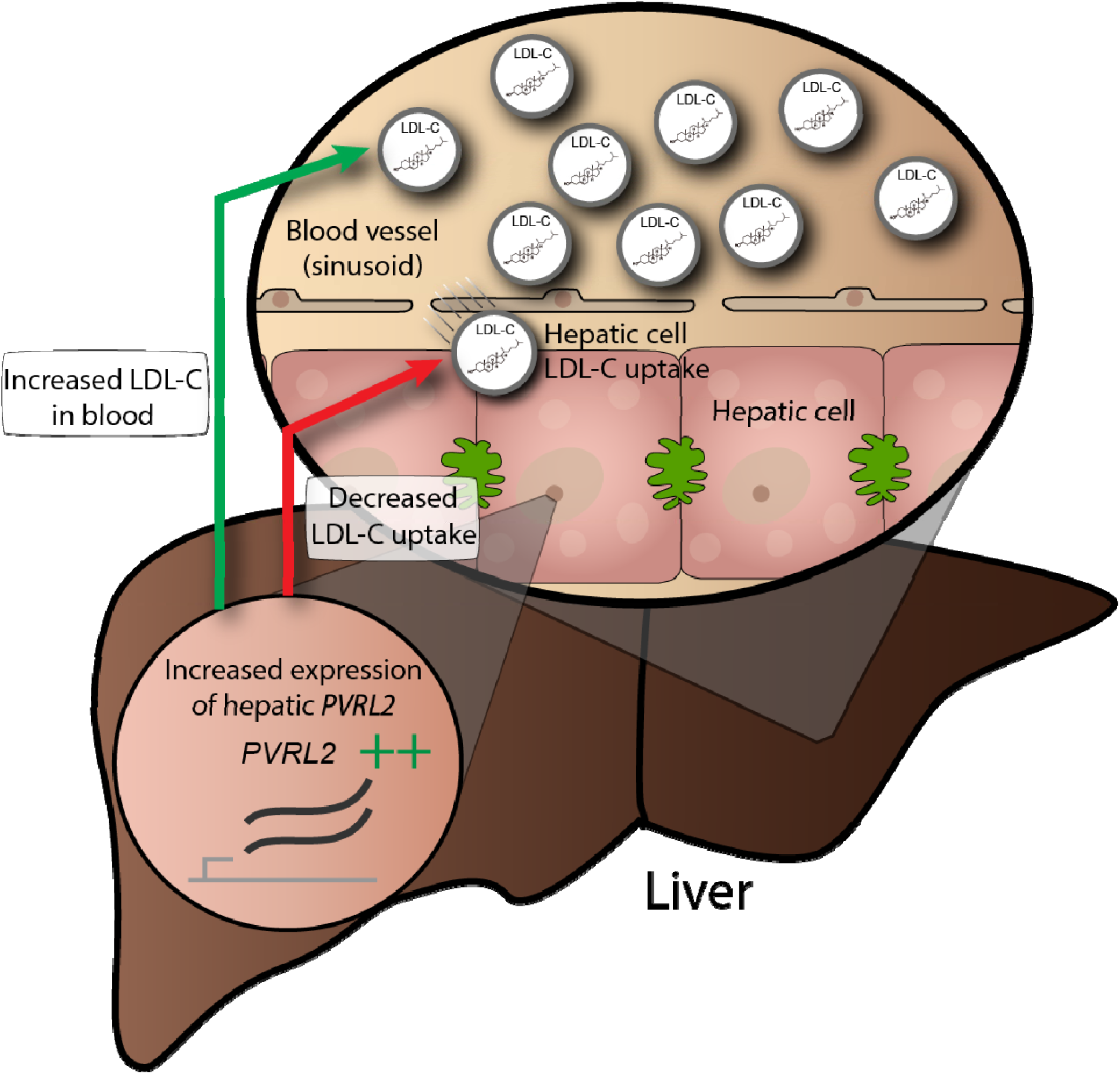
Biological interpretation of *PVRL2*. Functional and statistical evidence for the causal effect of *PVRL2* in liver on low density lipoprotein cholesterol (LDL-C) levels. The green arrow indicates a positive causal relationship between *PVRL2* and LDL-C levels in plasma – this relationship was detected in our analysis. The red arrow indicates a negative causal relationship between *PVRL2* and LDL-C uptake in hepatic cells – this relationship was detected in small interfering RNA (si) experiments described in Blattman et al^44^.

In the MR analysis using eQTLs from liver, all the genes identified fall within LDL-C GWAS loci. Among these, we found a negative causal effect for the well-known *SORT1* gene (*p* = 5.9×10^−9^). This gene encodes the protein *sortilin* that has been functionally proven to down-regulate lipoprotein metabolism and hepatic lipoprotein export in liver^31,38–40^ (**Table 3** and **Supplemental Table 5**). We also found two other genes in the same GWAS locus, *PSRC1* and *CELSR2*, but the IV (only one was found) for these genes was identical to that of *SORT1*, due to the high correlation between expression levels of these genes. Full overlap of one single IV in this locus makes it is impossible to discern causal from pleiotropic genes using MR-methods, including MR-link. The fourth gene found to be significant using liver eQTLs is *PVRL2* (*p* = 3×10^−14^), which is located in the *APOE* locus associated to LDL-C (**Table 3**)^31,32^. For *PVRL2*, we estimated a positive causal effect; higher expression of *PVRL2* is causally related to higher LDL-C (Table 3). *PVRL2* is 17.5kb downstream of the *APOE* gene, and three common missense polymorphisms in *APOE* account for a large fraction of the association signal^32^. Interestingly, in the most recent GWAS meta-analysis for lipids, 19 jointly significant LDL-C variants were found spanning a 162Kb region that encompasses *PVRL2*^*32*^. This indicates that, while *APOE* plays a major role, other genes in this locus are also likely involved in LDL-C regulation and that pleiotropic effects are to be expected. Our analyses indicate that *PVRL2* is one of the causal genes at this locus. The positive effect of *PVRL2* on LDL-C was also seen in the analysis that used blood eQTLs from BIOS (*p* = 4.3×10^−5^), although it did not pass our significance threshold (*p* = 4.6×10^−6^). Of note, since LD between IVs used in the analysis of blood and liver eQTLs was low (r^2^ < 0.2), the results potentially indicate a dual causal role for the gene across these two tissues.

*PVRL2* has mostly been studied in the context of atherosclerosis, where it has been shown to act as cholesterol-responsive gene involved in trans-endothelial migration of leukocytes in vascular endothelial cells, a key feature in atherosclerosis development^41–43^. Our results indicate a role for *PVRL2* in modulating plasma levels of LDL-C via its expression variation not only in blood but also in the liver. Biologically the role in liver could be explained by increased production of very-LDL or a decreased LDL-C uptake (**Figure 3**). In line with this hypothesis, a siRNA screen in hepatic cell lines of genes in the APOE locus showed that down-regulation of *PVRL2* gene expression promotes LDL-C uptake^44^ (**Figure 3**). Overall, our results and existing functional evidence support that *PVRL2* expression is correlated with LDL-C levels and show, for the first time, a causal effect in liver (**Figure 3**).

## Discussion

Identification of genes whose changes in expression are causally linked to a phenotype is crucial for understanding the mechanisms behind complex traits. While several methods exist that infer causal relationships between two phenotypes, these rely on a set of assumptions that are often violated when gene expression is the exposure. Specifically, the presence of LD and pleiotropy between the genetic variants chosen as IVs are the main cause of violations of such assumptions^17,18,20,45^. Here we interrogated a large gene-expression dataset and showed that the eQTLs of a gene, which can be used as IVs, are very likely to be in LD, but not overlapping, with eQTLs of other genes, indicating that potential sources of pleiotropy in transcriptome-wide MR analyses are likely from variants in LD with the IVs.

We therefore developed MR-link, a novel causal inference method that is robust to unobserved pleiotropy. Our *in-silico* results show that MR-link outperforms all the other MR methods tested and has well-controlled FPR and high statistical power. MR-link jointly models the outcome using jointly significant eQTLs as IVs, combined with variants in LD, to correct for all potential sources of pleiotropy. To our knowledge, this is the first time that this approach is used in a causal inference method.

We tested MR-link’s performance with real data by applying it to LDL-C cholesterol measurements and eQTLs derived from blood, cerebellum and liver. This identified known and novel key player genes within and outside GWAS loci. For example, in liver we identified the well-known negative causal relationship between expression of *SORT1* in liver and LDL-C^38–40^. In liver, and suggestively in blood, we detected a causal effect for *PVRL2*, a gene located in the *APOE* locus. While a role for this gene is mostly known for immune and endothelial cells and in the context of atherosclerosis^41–43^, our results indicate that regulation of expression of this gene in both blood and liver causally affects LDL-C levels. Given its established role in atherogenesis, *PVRL2* has been proposed as a potential therapeutic target for atherosclerosis. Our study indicates that such strategies should not only take into account the effect on atherosclerotic plaques but also consider the hepatic function of *PVRL2* in regulating plasma LDL-C levels in humans.

All the genes identified in the analyses that used eQTLs from blood were different from those identified using eQTLs from liver. While this is partly due to statistical power, as the BIOS cohort is more than 20 times larger than the GTEx cohort used to derive eQTLs in liver, this may also be related to tissue-specific functions. We expect that causal genes found in whole blood will affect LDL-C through pathways that signal for lipid changes, whereas genes found in liver are more likely to be involved in lipid metabolism.

MR-link has several advantages over other recent MR methods developed to overcome bias from LD and pleiotropy^17,23^. First, MR-link can model unobserved pleiotropy, whereas sources of pleiotropy need to be specified in multivariate MR methods. This is particularly important because sources of pleiotropy may be context-dependent and may arise from a phenotype other than those being measured in a cohort^14,30^. Second, MR-link can derive robust causal estimates even when only one or two IVs are available. The majority of genes tested in our large eQTL dataset have fewer than three IVs (68%), which makes it impossible for MR-PRESSO, MR-Egger and LDA-MR-Egger to make causal estimates^17,18,20^.

The application of MR-link is not restricted to gene expression; it can also be applied to other molecular layers that are known to have a similar genetic architecture to gene expression, such as proteins or metabolites. Given the increases in sharing of summary statistics from functional genomics QTL studies, coupled with the development of very large biobanks such as the UK biobank, the Estonian Biobank, the Lifelines cohort study and the Million Veteran Program cohort^46–49^, we foresee many opportunities for applications of MR-link to individual-level data for the identification of the molecular mechanisms underlying complex traits. Of note, while we have limited our simulations to quantitative traits as an outcome in this paper, MR-link could be applied to binary traits such as human diseases. However, we have not investigated its performance in detail for binary outcome phenotypes. Furthermore, as for all MR studies, our method can be applied to populations of any ethnicity, provided that the exposure summary statistics are derived from a population that is ethnically-matched with the outcome cohort.

We foresee that many causal relationships will be discovered as highly powered causal inference methods such as MR-link are applied to many human traits. This could make it possible to build extensive causal networks similar in size and complexity to metabolic networks of small molecules, which would provide valuable insights into the mechanisms behind human traits and diseases.

### Data availability

Individual level data of BIOS and Lifelines cohorts are available upon request to their respective biobanks (https://www.bbmri.nl/acquisition-use-analyze/bios and https://www.lifelines.nl/researcher). GTEx summary statistics can be downloaded from the GTEx website (https://gtexportal.org/home/datasets/)

### Code availability

An implementation of MR-link, the methods to recreate the simulated data and instructions on usage can be found at https://github.com/adriaan-vd-graaf/genome_integration.

## Acknowledgments

We are very grateful for the altruistic donation of biological materials by our generous study participants, without them this study would not be possible. In addition, we thank the UMCG Genomics Coordination center, the UG Center for Information Technology and their sponsors BBMRI-NL & TarGet for storage and computing infrastructure. We thank BBMRI-NL for providing the transcriptome and genotyped data for the BIOS cohort. We thank P. Visscher, N. Wray, J. Yang, E. Lopera-Maya, O. Bakker and N. de Klein for valuable advice during the development and writing of this work. We thank K. McIntyre for editorial assistance. This work is financed by the Netherlands Organization for Scientific Research (NWO): NWO Spinoza Prize SPI 92-266 to C.W., and by Fondation Lefoulon-Delalande (to A.R.).

## Author contributions

A.v.d.G. conceived and designed the MR-link framework with critical input from S.S.; A.v.d.G. performed all data analyses on the datasets used in this study; A.v.d.G. and A.C. performed quality control analyses on the BIOS and Lifelines cohorts; B.C. and C.W. provided access to the datasets used in this study; A.v.d.G. and S.S. wrote the manuscript with critical inputs from Y.L., H.J.W., A.R. and A.C.; Y.L., S.S. and C.W. supervised the study; C.W. provided funding for this study. All authors read and approved the manuscript.

## Methods

### BIOS consortium cohort genotype and expression analysis

We used genotype and expression measurements on 3,746 Dutch individuals from the Biobank-based Integrative Omics Study (BIOS; http://www.bbmri.nl/acquisition-use-analyze/bios/), a collection of six different data cohorts: Lifelines DEEP^29^, Prospective ALS Study Netherlands^58^, Leiden Longevity Study^59^, Netherlands Twin Registry^60^, The Cohort on Diabetes and Atherosclerosis Maastricht^61^ and the Rotterdam Study^62^. Genotyping was performed separately per cohort (see references). All combined genotypes were imputed to the Haplotype reference consortium dataset^63^ using the Michigan imputation server^64^. We retained only biallelic SNPs and confined our analyses to variants with minor allele frequency (MAF) > 0.01, Hardy-Weinberg equilibrium (HWE) p value < 10^−6^ and an imputation imputation server^64^. We retained only biallelic SNPs and confined our analyses to variants with minor quality RSQR > 0.8. A genetic relationship matrix (GRM) was derived based on LD-pruned genotypes using the Plink 1.9 command “--indep 50 5 2”, and one individual was kept from all pairs of individuals that had a GRM value > 0.1 using the “--rel-cutoff” Plink 1.9 command^28^. Population outliers were identified using a principal component analysis of the GRM, and individuals more distant than three standard deviations from the mean of principal component 1 and principal component 2 were removed.

RNA-seq gene expression quality control and processing has been described previously^13^. In brief, RNA extracted from whole blood was paired end sequenced using the Illumina HiSeq 2000 instrument. RNA-seq read alignment was performed using STAR (version 2.3.0e)^65^. During alignment, variants with MAF < 0.01 from the Genome of the Netherlands were masked^66^. Gene expression was quantified using HTSeq^67^. Samples with less than 80% of reads mapping to exons were considered of low quality and removed. Samples were also removed if they had less than 85% of mapped reads, or if they had a median 3’ bias larger than 70% or smaller than 45%. To further account for unobserved confounders, the expression matrix was corrected for the first 25 principal components as well as 5’ bias, 3’ bias, GC content, intron base-pair percentage and sex following the procedure of Zhernakova et al.^13^. After genotype and expression quality control filters, 3,503 individuals with expression data of 19,960 transcripts and genotype information of 7,838,327 SNPs were available for analyses. eQTL association analysis was performed for SNPs located ±1.5 Mb of the transcript using plink 1.9 and the ‘--assoc’ command^28^. For 10,831 genes at least one eQTL at p < 5×10^−8^ was identified, and those genes were used for all the analyses described in this manuscript.

We quantified how many genetic variants are necessary to explain gene expression using a conditional joint analysis approach. We identified jointly significant eQTLs by applying GCTA-COJO^25^ to eQTL summary statistics, using the BIOS cohort as LD reference panel, and selecting jointly significant variants that showed a p < 5 × 10^−8^ in this analysis step. To infer how often eQTLs are shared between genes, we assessed the percentage of genes with top eQTLs (or jointly significant variants) that have LD r^2^ > 0.5.

### Lifelines cohort genotype data and low-density lipoprotein cholesterol levels

Lifelines is a multi-generational cohort study of 167,000 individuals from the north of the Netherlands. It was approved by the medical ethics committee of the University Medical Center Groningen and conducted in accordance to the Helsinki Declaration Guidelines. All participants signed an informed consent form prior enrollment. A subset of 13,283 samples were genotyped with cytoSNP array and underwent quality control as previously described^49^. After genotype quality control, samples were imputed using the Genome of the Netherlands reference panel^66^ and Minimac version 2012.10.3^49,68^. Variants were excluded if they were of bad imputation quality (RSQR < 0.3), showed deviation from HWE (p < 10^−6^), or if they were absent in the set of quality controlled genotyped and imputed variants of the BIOS cohort.

Low-density lipoprotein cholesterol (LDL-C) was estimated using the Friedewald equation^69^, based on triglycerides, high density lipoprotein and total cholesterol levels^49^. Total cholesterol levels of individuals who were prescribed cholesterol-lowering medication were divided by 0.8 prior calculating LDL-C. Individuals with >4.52 mmol/Liter total triglycerides were removed^69^. Additionally, LDL-C levels were corrected for age, age squared and sex. After genotype and LDL-C quality control, 12,449 individuals and 7,336,374 variants remained for analyses. Association analysis for additive effects on LDL-C was performed using linear regression on standardized genotypes, e.g. transforming genotypes into a distribution with mean 0 and variance 1. Summary statistics of this analysis were used to perform MR analyses using existing MR methods listed in Table 1.

### GTEx download and analysis

We downloaded GTEx version 7 eQTL summary statistics, including non-significant results, from the GTEx website (https://gtexportal.org/home/datasets/)^24^. For every gene with at least one eQTL at *p* < 5×10^−8^, conditional analysis using GCTA-COJO was performed to select secondary variants at the same threshold, using the BIOS cohort as an LD reference. This resulted in in 4,028, 1,557 and 1,726 genes with at least one jointly significant eQTL for whole blood, liver and brain (cerebellum) tissues, respectively.

### Simulation of genotypes

403 non-Finnish European individuals were isolated from the 1000 Genomes phase 3 release and used as a starting point for genotype simulation^27^. We simulated genotype data for 25,000 individuals in a chromosomal region (Chromosome 2, 100Mb to 105Mb, human genome build 37) using the HAPGEN2 program, combined with interpolated HAPMAP3 recombination rates^28,70^. The region was then reduced to 1Mb in length: between 102Mbp and 103Mb. Only biallelic SNPs with MAF < 0.01 were retained from simulated genotypes, leaving 3,101 variants in this region. Simulated individuals were separated into an outcome cohort of 15,000 individuals, and into an exposure cohort and an LD reference cohort of 5,000 individuals each. These cohort sizes were chosen to roughly represent the sizes of BIOS and Lifelines cohorts.

### Simulation of phenotypes

We simulated quantitative phenotypes representing the exposures by randomly selecting SNPs from the simulated genetic region, and subsequently assigning these an effect. Causal SNPs were selected to represent both pleiotropy through LD (**Figure 1C**) and pleiotropy through overlap (**Figure 1B**). For the scenario of pleiotropy through LD (**Figure 1C**), one to ten causal SNPs were randomly selected to be causal to the exposure, and the same number of SNPs was selected among all SNPs in moderate LD (0.25 < r^2^ < 0.95) for being causal to the unobserved (pleiotropic) exposure.

When pleiotropy through overlap was simulated (**Figure 1C**), the causal SNPs for the observed and unknown exposure were selected to be identical. A combination of pleiotropy through overlap and pleiotropy through linkage was simulated by choosing some or all of the SNPs of the unobserved exposure to be overlapping and some being in LD (0.25 < r^2^ < 0.95).

The mathematical framework is as follows. For each selected causal SNP of the exposure (subset *S*_*E*_), we simulated an effect-size from the uniform distribution *U (*– 0.5,0.5), and then simulated the observed exposure as:

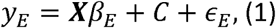

where ***X*** is a genotype matrix of size *n × m*, with *n* being the number of individuals (5,000) and *m* the number of variants in the region (3,101 in the simulated data), *β*_*E*_ is the vector of effects 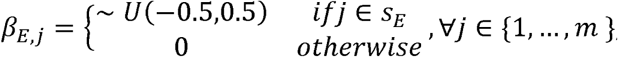 and *C* ∼ *N*(0,0.5)^*n*^ is a matrix of specific confounder per individual. Finally, *ϵ*,_*E*_∼*N*(0,1) ^*n*^ is the measurement error of the exposure. Similarly, the unobserved exposure *y*_*U*_ was simulated as:

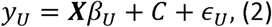

where, *b*_*U*_ is the vector of effects defined as: 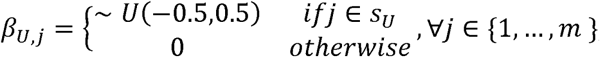, *S*_*U*_ is the selection of SNPs for the unobserved exposure and *ϵ*_*U*_ are measurement errors distributed as *ϵ*_*E*_. The outcome phenotype. *y*_*o*_ was then simulated as a linear combination of the observed and unobserved exposures:

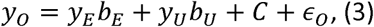

where the causal effect of interest is *b*_*E*_ and the (unknown) pleiotropic effect is *b*_*U*_. Again, the measurement error *ϵ*_*o*_ is drawn from the standard normal distribution.

The effects *β*_*E*_, *β*_*U*_, *b* _*E*_ and *b*_*U*_ were randomly drawn from their respective distribution and used in both cohorts (exposure and outcome), while the other random variables were randomly drawn in a cohort-specific manner. Since our model was built to account for unobserved pleiotropy, only the outcome phenotypes and the summary statistics of the (observed) exposure phenotype were used in the causal inference analysis.

### Simulation parameters and scenarios

We simulated 1,500 runs per scenario, each with a unique outcome (*O*) and two exposures (*E* and *U*). The scenarios differed in the number of causal SNPs (which varied from one to ten for both the observed and unobserved exposure); the strength of the causal relationship of interest (varied from no causal effect up to a large effect (*b*_E_ ∈ 0,0.05,0.1,0.2,0.4); and the presence (*b*_*U*_ = 0.4) or absence (*b*_*U*_ = 0.0) of the pleiotropic effect. This resulted in 10*5*2 = 100 different scenarios.

In certain cases, an estimate cannot be made by an MR method, for instance when insufficient IVs are identified or a solution is not found in the estimation method. As a result, there are sometimes fewer estimates than expected in the final results. To ensure the stability of our FPR and power estimates, we have only reported results for a MR method in a specific scenario if we had more than 100 estimates out of the 1,500 simulated runs.

### Instrumental variable selection

IV selection can be difficult when there is LD between association signals. In simulations, we used two IV selection techniques: GCTA COJO^25^ and p value clumping, using standard settings of plink 1.9 except for the r^2^ threshold, which was set to 0.1^28^. Both selection methods used a *p* value threshold of *p* < 5×10^−8^. When selecting IVs for BIOS and GTEX, we only used the GCTA-COJO technique.

### MR-link

MR-link is a method for causal inference that is robust to the presence of LD and unknown pleiotropy. It is a two-sample MR approach that requires individual-level data from the outcome cohort and summary statistics (effect sizes, standard errors and minor allele frequencies) from an exposure. Conceptually, MR-link jointly models a known exposure with SNPs that are in LD with the exposure IVs (tag-SNPs). Tag-SNPs are used to account for the unobserved pleiotropic effect present in a locus.

We defined our model in the following manner. Let ***x*** be a genotype matrix of *n* × *m*, where *n* is the number of individuals in the outcome study and *m* are all the SNPs in a *cis*-region around the transcript (±1.5 Mb of the transcript), in which SNPs at indices *S*_*E*_ are the causal genetic variants (IVs) for the exposure *E*. If we define the exposure, and the unobserved (pleiotropic) exposure *E* and the unobserved (pleiotropic) exposure *U* as in equation (10 and (2), then the outcome phenotype *y*._*o*_ from equation (3) can be represented as a function of *E*, and *U* with the following equation:

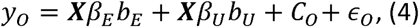

where *b*_*E*_ is the causal effect of interest of the exposure on the outcome, *b*_*U*_ is the causal effect of the unobserved exposure, *C*_*0*_ represents some confounder and ϵ_0_ is the measurement error of the outcome In the case of pleiotropy through overlap, all variants will be the same, thus *S*_*E*_*=S*_*U*_ In the hypothetical case that the genetic effects for both the exposure *E* and the pleiotropic exposure *U* are known, we can estimate (*b*_*E*_ by solving equation (4). In a real-case scenario, only the IV(s) for the exposure are known, while the variants that contribute to the unknown (pleiotropic) exposure anexposure are known, while the variants that contribute to the unknown (pleiotropic) exposure and their effect on the outcome are unknown.

MR-link uses the following procedure to estimate causal effects:

1. After selection of 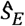 IVs for the exposure, conditional effect sizes 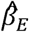 are determined for these IVs using the GCTA-COJO method^25^. A vector of effect sizes” 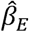 for all SNPs in the region is thus defined as: 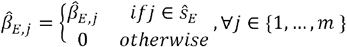
2. All SNPs in LD 0.1< r^2^< 0.99 with the exposure IVs are potential tag-SNPs. These variants are iteratively pruned for high LD so that tag-SNPs, *S_T_*, are always r^2^ <0.95 with each other in order to reduce collinearity and computation time.
3. The following equation is solved for *b*_*E*_ using ridge regression:

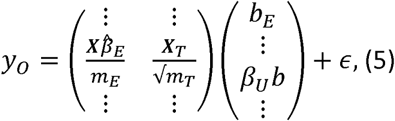

where ***x***_*T*_ genotype matrix of the outcome containing only tagging variants as defined in (2) and *m*_*T*_ is the number of tagging variants and *m*_*E*_are the number of IVs selected by the selection method.

We also considered solving the equation (5) using ordinary least squares. However, due to the multicollinear nature of the 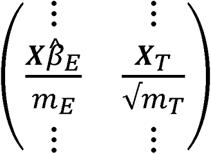 matrix, this approach leads to very low detection power (**Supplemental Table 2, Supplemental Figure 4, 5, 6** and **Supplementary Note**). We therefore applied ridge regression to solve the equation and determined a T statistic and subsequent Wald test *p* value for ridge regression^71^. Due to the over-conservative nature of the resulting p value in simulations and real data (**Supplemental Table 2, Supplementary Figure 4, 5, 6** and **Supplementary Note**), we adjusted the p value distribution of each different scenario by fitting a beta distribution to all estimates to calibrate the final *p* values (Supplementary Note). When we report results for MR-link in simulated data, it is these adjusted *p* values that we are referring to.

Since sparsely genotyped regions and/or highly independent IVs may affect MR-link performance (increased FPR) due to the reduced number of tag-SNPs that explain a residual outcome signal, we applied a permutation procedure to assess the robustness of significant results in applications to real data. Specifically, for all genes that passed the Bonferroni significance threshold, we permuted the outcome phenotypes (LDL-cholesterol in our case) 1000 times, recalculated causal estimates with MR-link, and re-classified the gene as non-significant if more than 1% of permutations had a more significant causal effect than originally observed.

### Mendelian Randomization analyses

Causal relationships were estimated with MR-link and four other existing methods: Inverse variance weighting (IVW)^45^, LDA-MR-Egger regression^17^, MR-Egger regression^18^ and MR-PRESSO. All methods were (re-)implemented in Python and compared to present equal results when compared with their original implementation. The corresponding code is available https://github.com/adriaan-vd-graaf/genome_integration.

The IVW method is a weighted meta-analysis of causal estimates from single IVs. Specifically, a causal estimate *b*_*i*_,for an IV *i* is estimated 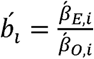 where 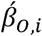 is the marginal effect of SNP*i* on the outcome and” 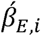 is the marginal effect of the exposure. For the estimation of the causal effect, single IV causal estimates are combined using weights proportional to the inverse variance of such estimates using the two-terms definition of standard error: 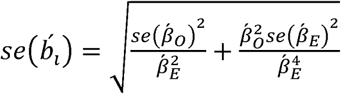 as described in^72^.

MR-Egger regression adjusts for average pleiotropy by fitting a linear regression between the exposure SNP-effects and the outcome SNP-effects^18^. It assumes that <50% of the variants have a pleiotropic effect. MR-Egger can be applied when three or more instruments are available.

LDA-MR-Egger is similar to MR-Egger but also recognizes LD. LDA-MR-Egger can only be used when LD information between the IVs is available^17,19^.

MR-PRESSO is a method of causal inference that implements an approach to identify and remove outliers from the IVW framework^20^. It assumes that <50% of the variants have a pleiotropic effect. MR-PRESSO is unable to adjust for the presence of pleiotropy if fewer than three IVs are available, of if fewer than two IVs are left after outlier correction.

For these four methods we used LDL-C full GWAS summary statistics derived from the association carried out in the Lifelines study, as described above.

