## Supplemental Notes for "A novel Mendelian randomization method identifies causal relationships between gene expression and low-density lipoprotein cholesterol levels"

### Supplemental materials

**Supplementary Note.**

**Supplemental Figure 1.** Simulation results depicting Type I error rates (at 0.05 significance) for two different IVs selection methods

**Supplemental Figure 2.** Comparison of IVs and effect-sizes identified when using eQTLs from BIOS and GTEx whole blood

**Supplemental Figure 3**. Forest plots of causal effect estimates for the 15 significant genes identified by MR-link using BIOS blood eQTLs

**Supplemental Figure 4.** Simulation results using different solvers for MR-link

**Supplemental Figure 5.** Calibration of *p* values in null scenarios

**Supplemental Figure 6.** Comparison of MR-link p values obtained using the BIOS eQTLs data and LDL-C individual data

**Supplemental Table 1.** False positive rates and power of different MR methods and IV selection when no pleiotropy is simulated

**Supplemental Table 2.** False positive rate and power of different MR methods when pleiotropy through linkage disequilibrium is simulated

**Supplemental Table 3.** False positive rate and power of different MR methods when pleiotropy through overlap is simulated

**Supplemental Table 4.** Full results of the genes that were significant using MR-link and the eQTL from the BIOS cohort.

**Supplemental Table 5.** Significant genes identified in the MR-link analysis that used GTEx eQTL data

**Supplemental Table 6.** Results of all other MR methods tested for the 15 genes identified using MR-link and the BIOS cohort.

### Supplementary Note

#### Calibration of *p* values resulting from MR link analysis

In a preliminary version of MR-link we used Ordinary Least Squares (OLS) to solve equation (5) (**Methods**) (MR-link, OLS). Here we observed that MR-link with OLS implementation had FPR close to expectations (**Supplemental Figure 4**) (**Supplemental Table 2**), but lacked sufficient power to detect a causal effect (power ranged from 0.11 to 0.19 for the 10 different scenarios). We hypothesized that this was likely due to multicollinearity of the model. To overcome this issue, we replaced OLS with ridge regression, which is relatively robust to multicollinearity compared to OLS and LASSO regression, especially when $n < m$ as is the often the case in our analyses[1,2]. We used a previously described[3] method to estimate the standard error of the ridge estimate and subsequently derived a T statistic. However, using MR-link with ridge regression implementation was very conservative (**Supplemental Table 2-4, Supplemental Figure 4** and **5**). To ensure that the MR-link ridge test statistic *p* value followed a uniform distribution in the case of the null scenario, we adjusted the *p* values using a beta distribution fit. The beta distribution estimate was made using a Markov Chain Monte Carlo approach, with the parameters of the beta distribution having a uniform prior between 0 and 20. These parameters are estimated using PYMC3 combined with a No-U-Turn sampler[4]. This estimation scheme for the beta distribution parameters is preferable to maximum likelihood (ML) estimates because ML estimates are prone to convergence issues when a distribution is heavily shifted to one.

In simulations, the beta distribution is estimated in the null scenario (no causal effect), then applied to all scenarios. The results shown in **Supplemental Figure 4** and **Supplemental Figure 5** demonstrate that calibration successfully restored *p* values to follow the expected distribution.

In the application to real data, the estimation of the beta distribution for calibration was done on all observations assuming that the majority of tests in a transcriptome-wide analysis represent a null causal effect. This concept is similar to that behind the genomic control correction approach used in GWAS studies[5]. When we applied this approach to the results obtained using eQTLs from BIOS and LifeLines individual data, we saw a similar behavior to that observed in simulations: *p* values obtained using MR-link with OLS implementation follow the expected uniform distribution (**Supplemental Figure 6A**), whereas those obtained using *MR-link* with ridge regression implementation indicate that the test is conservative (**Supplemental Figure 6B**), and those obtained using MR-link with ridge regression and *p* value calibrations follow the expected uniform distribution before deviating from the expectation to represent a limited number of genes with a true causal effects (**Supplemental Figure 6B**). All the results presented in the main text for simulations and application to real data refer to MR-link with ridge regression implementation and calibration of *p* values. For application to real data, we also provided permutation results of calibrated *p* values.

**BIOS Consortium Members**

*Management Team*

Bastiaan T. Heijmans (chair)[1], Peter A.C. 't Hoen[2], Joyce van Meurs[3], Rick Jansen[5], Lude Franke[6].

*Cohort collection*

Dorret I. Boomsma[7], René Pool[7], Jenny van Dongen[7], Jouke J. Hottenga[7] (Netherlands Twin Register); Marleen MJ van Greevenbroek[8], Coen D.A. Stehouwer[8], Carla J.H. van der Kallen[8], Casper G. Schalkwijk[8] (Cohort study on Diabetes and Atherosclerosis Maastricht); Cisca Wijmenga[6], Lude Franke[6], Sasha Zhernakova[6], Ettje F. Tigchelaar[6] (LifeLines Deep); P. Eline Slagboom[1], Marian Beekman[1], Joris Deelen[1], Diana van Heemst[9] (Leiden Longevity Study); Jan H. Veldink[10], Leonard H. van den Berg[10] (Prospective ALS Study Netherlands); Cornelia M. van Duijn[4], Bert A. Hofman[11], Aaron Isaacs[4], André G. Uitterlinden[3] (Rotterdam Study).

*Data Generation*

Joyce van Meurs (Chair)[3], P. Mila Jhamai[3], Michael Verbiest[3], H. Eka D. Suchiman[1], Marijn Verkerk[3], Ruud van der Breggen[1], Jeroen van Rooij[3], Nico Lakenberg[1].

Data management and computational infrastructure

Hailiang Mei (Chair)[1][2], Maarten van Iterson[1], Michiel van Galen[2], Jan Bot[1][3], Dasha V. Zhernakova[6], Rick Jansen[5], Peter van ’t Hof[1][2], Patrick Deelen[6], Irene Nooren[1][3], Peter A.C. ’t Hoen[2], Bastiaan T. Heijmans[1], Matthijs Moed[1].

*Data Analysis Group*

Lude Franke (Co-Chair)[6], Martijn Vermaat[2], Dasha V. Zhernakova[6], René Luijk[1], Marc Jan Bonder[6], Maarten van Iterson[1], Patrick Deelen[6], Freerk van Dijk[1][4], Michiel van Galen[2], Wibowo Arindrarto[1][2], Szymon M. Kielbasa[1][5], Morris A. Swertz[1][4], Erik. W van Zwet[1][5], Rick Jansen[5], Peter-Bram ’t Hoen (Co-Chair)[2], Bastiaan T. Heijmans (Co-Chair)[1].

[1] Molecular Epidemiology, Department of Biomedical Data Sciences, Leiden University Medical Center, Leiden, The Netherlands

[2] Department of Human Genetics, Leiden University Medical Center, Leiden, The Netherlands

[3] Department of Internal Medicine, ErasmusMC, Rotterdam, The Netherlands

[4] Department of Genetic Epidemiology, ErasmusMC, Rotterdam, The Netherlands

[5] Department of Psychiatry, VU University Medical Center, Neuroscience Campus Amsterdam, Amsterdam, The Netherlands

[6] Department of Genetics, University of Groningen, University Medical Centre Groningen, Groningen, The Netherlands

[7] Department of Biological Psychology, VU University Amsterdam, Neuroscience Campus Amsterdam, Amsterdam, The Netherlands

[8] Department of Internal Medicine and School for Cardiovascular Diseases (CARIM), Maastricht University Medical Center, Maastricht, The Netherlands

[9] Department of Gerontology and Geriatrics, Leiden University Medical Center, Leiden, The Netherlands

[10] Department of Neurology, Brain Center Rudolf Magnus, University Medical Center Utrecht, Utrecht, The Netherlands

[11] Department of Epidemiology, ErasmusMC, Rotterdam, The Netherlands

[12] Sequence Analysis Support Core, Department of Biomedical Data Sciences, Leiden University Medical Center, Leiden, The Netherlands

[13] SURFsara, Amsterdam, the Netherlands

[14] Genomics Coordination Center, University Medical Center Groningen, University of Groningen, Groningen, the Netherlands

[15] Medical Statistics, Department of Biomedical Data Sciences, Leiden University Medical Center, Leiden, The Netherlands

### Supplemental Figures

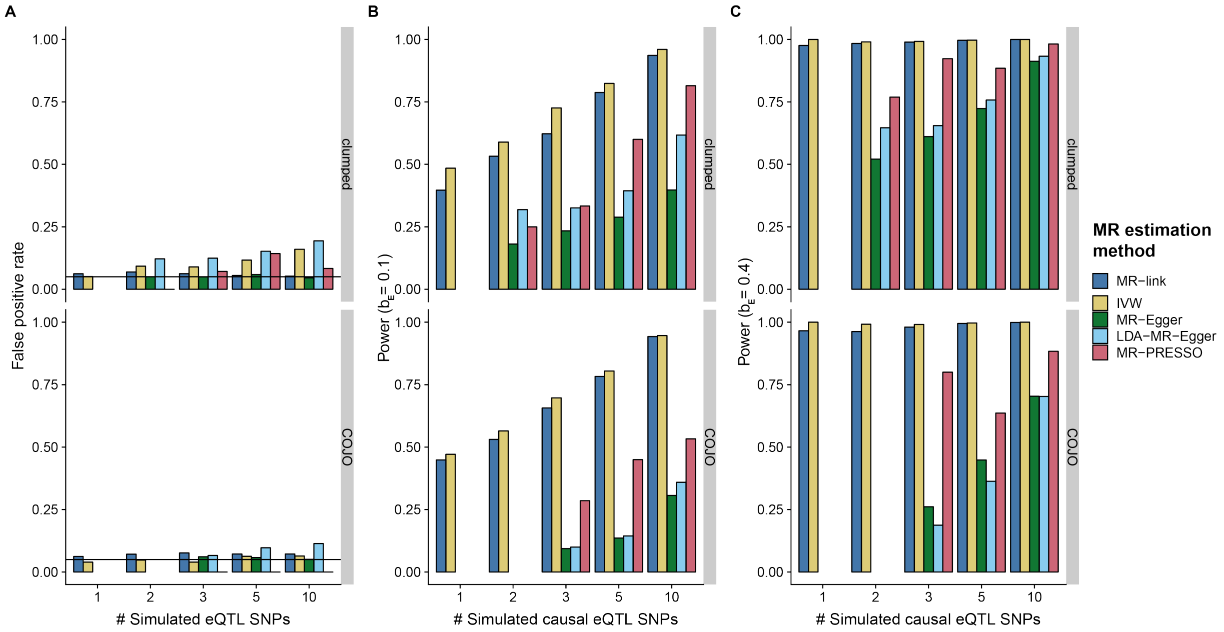


**Supplemental Figure 1. Simulation results depicting Type I error rates (at 0.05 significance) for two different IV selection methods**

Type I error rates (at 0.05 significance) in simulations when two different IV selection methods were used, GCTA-COJO (COJO) and p value clumping (clumped) in a non-pleiotropic scenario ($b_U = 0$) (**Methods**), with both selecting IVs at a threshold of p < 5x10$^{-8}$. (**A**) False positive rates in a scenario where no causal relationship is simulated. (**B**) Power to detect an effect in a scenario where a small causal effect is simulated. (**C**) Power to detect an effect in a scenario where a large causal effect is simulated. Note that MR-link and IVW are the only MR methods that can derive a causal estimate when only one or two instrumental variables are available.

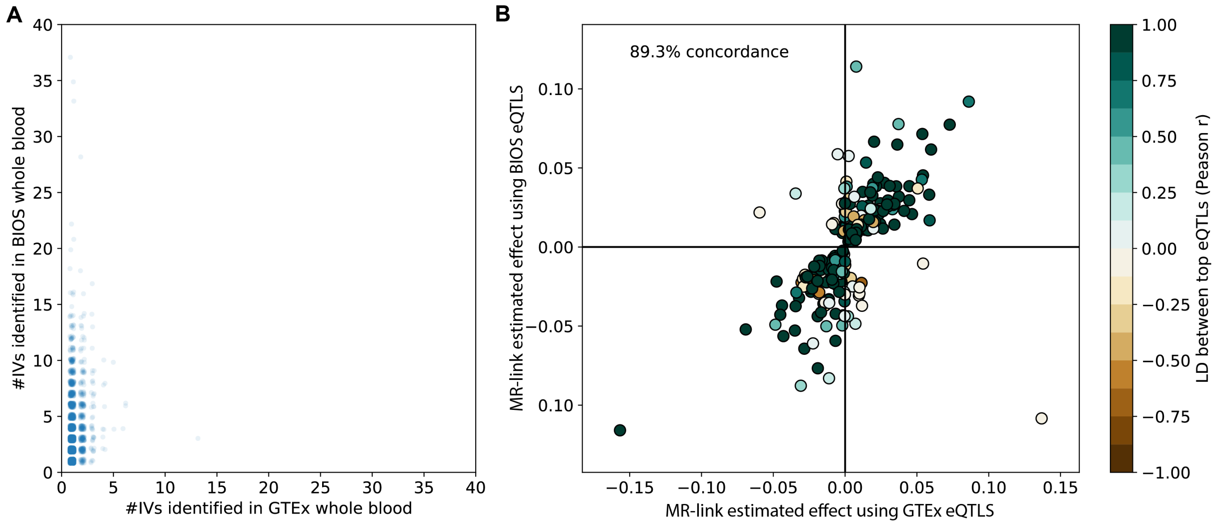


**Supplemental Figure 2. Comparison of IVs and effect-sizes identified when using eQTLs from BIOS and GTEx whole blood**

(**A**) The number of conditionally independent eQTL (IVs) identified for all the genes with at least one eQTL ($p < 5\text{x}10^{-8}$) in both the BIOS and GTEx cohorts. To represent the number of overlapping datapoints, all points are randomly jittered in both the x and y direction. (**B**) Scatterplot of causal effect sizes identified by MR-link when using GTEx whole blood and BIOS eQTLs for all the marginally significant ($p < 0.05$) genes detected using the BIOS cohort. Colors indicate the Pearson $r$ correlation between the most significant eQTLs of the gene.

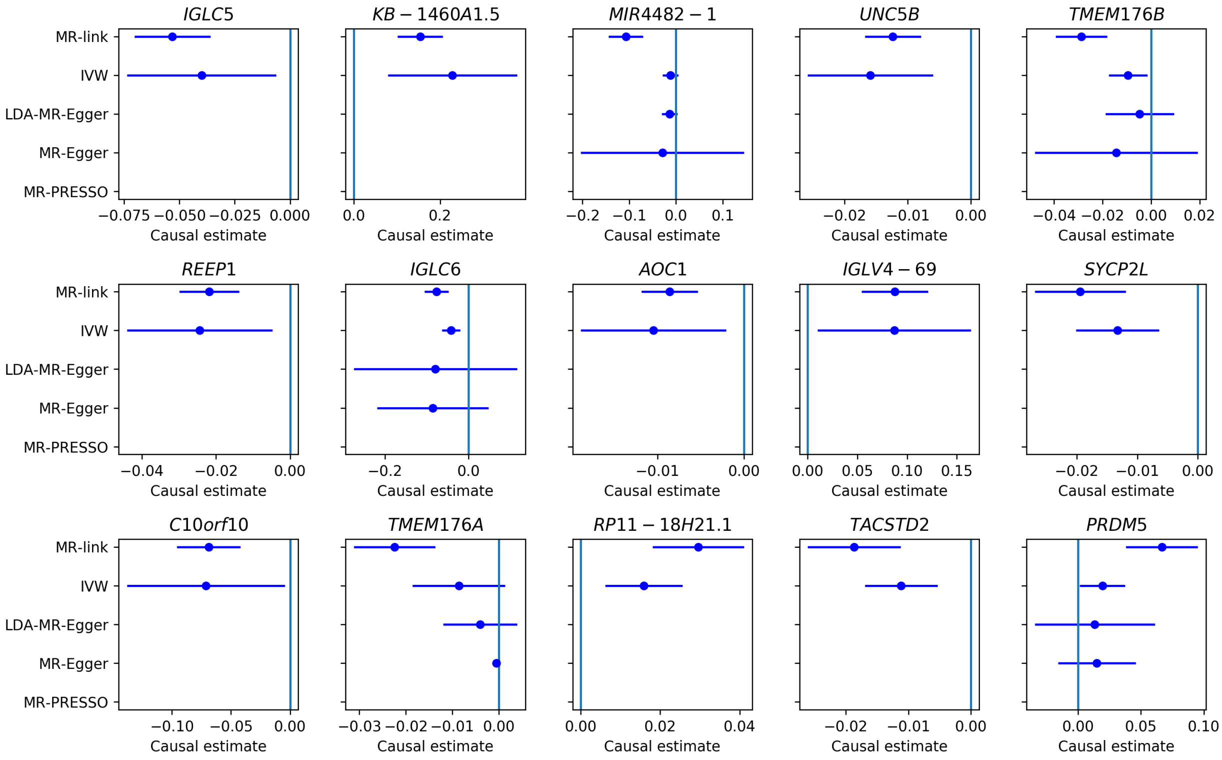


**Supplemental Figure 3. Forest plots of causal effect estimates for the 15 significant genes identified by MR-link using BIOS blood eQTLs**

Each plot displays the estimated causal effect size (X-axis) and 95% confidence interval (Y-axis) from all MR methods. No estimates for MR-PRESSO are shown because it identified too many outliers in all 15 genes identified by MR-link.

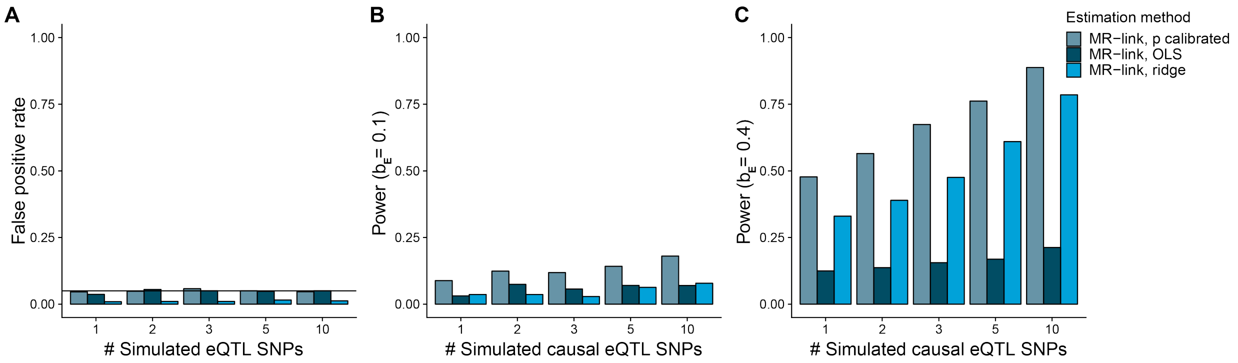


**Supplemental Figure 4. Simulation results using different solvers for MR-link**

Simulation results depicting false positive rates and detection power (at 0.05 significance) for MR-link when different equation solvers are used: OLS, ridge regression, and ridge regression followed by *p* value calibration (**Supplementary Note**). Results are from the same pleiotropic scenarios, as simulated in **Figure 2D-F.** (**A**) False positive rates in a scenario where no causal relationship is simulated. (**B**) Power to detect an effect in a scenario where a small causal effect is simulated. (**C**) Power to detect an effect in a scenario where a large causal effect is simulated.

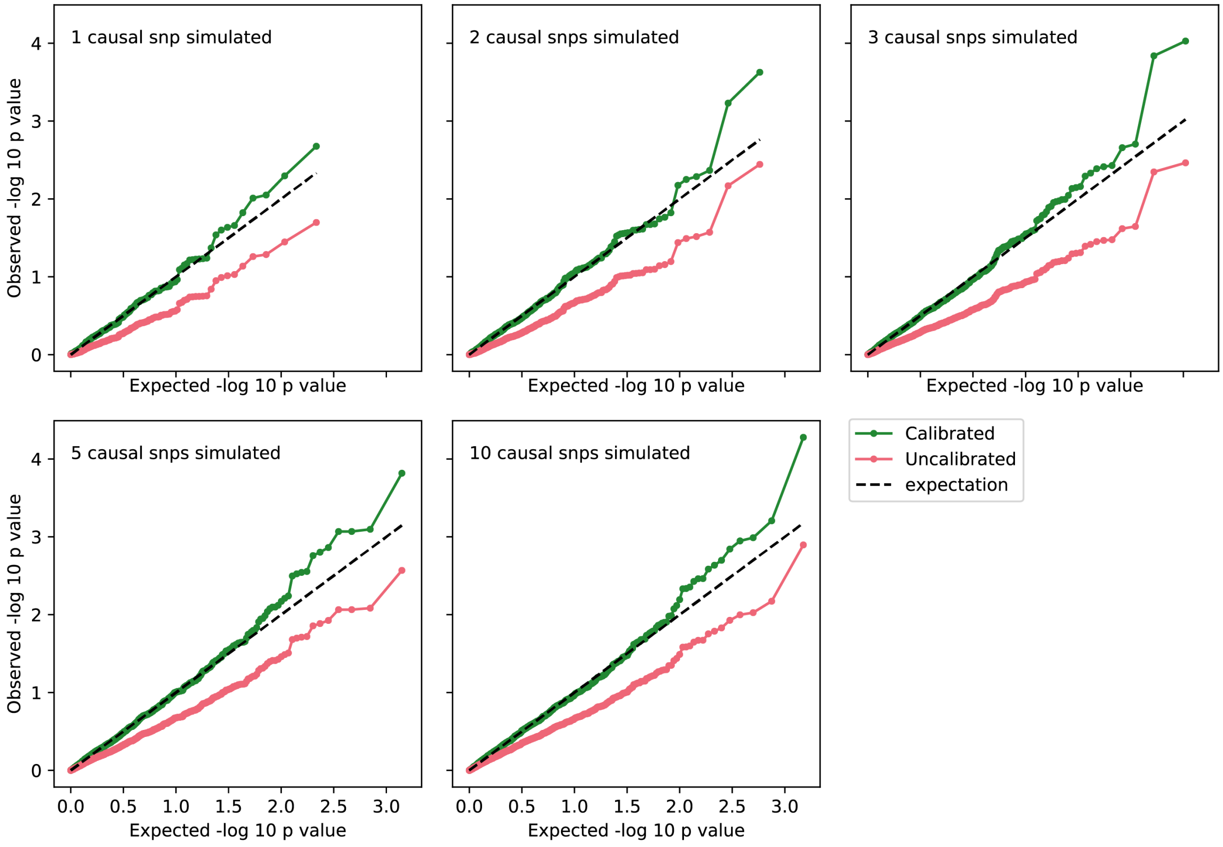


**Supplemental Figure 5. Calibration of *p* values in null scenarios**

Quantile-quantile plot of the p values obtained with MR-link ridge before and after calibration (beta distribution fit, see **Supplementary Note**) in the simulation scenario of pleiotropy and linkage with different numbers of causal SNPs (depicted in **Figure 2D**). Expected *p* values are calculated as the inverse of the rank of the *p* value sorted by significance. Dashed line is the diagonal.

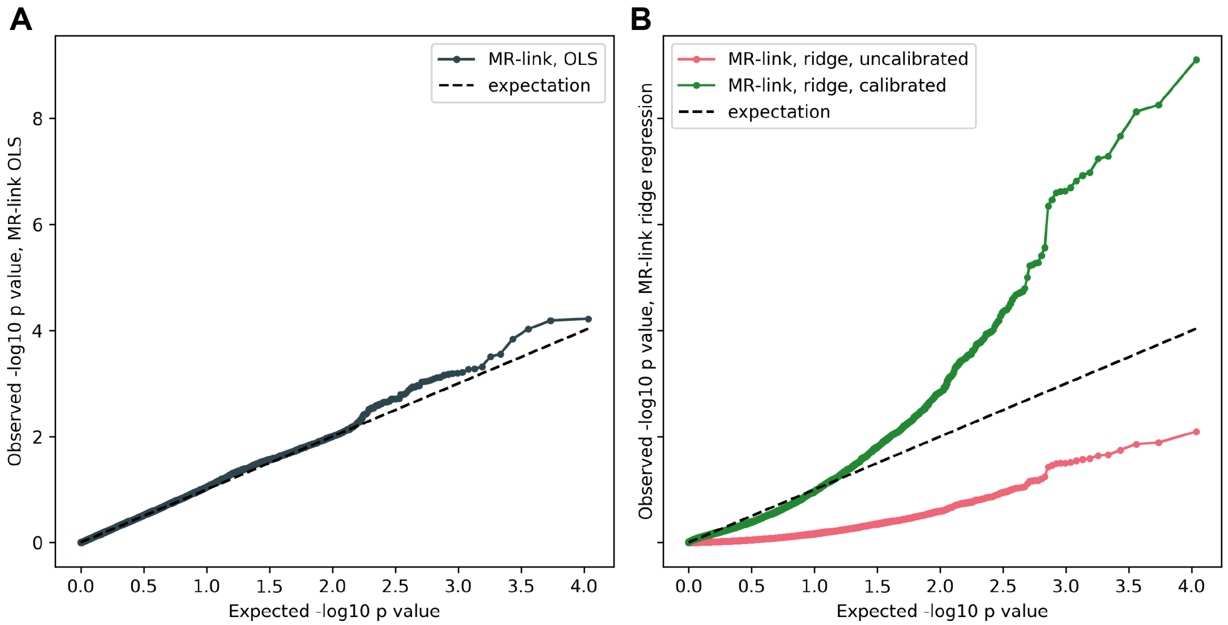


**Supplemental Figure 6. Comparison of MR-link *p* values obtained using the BIOS eQTLs data and LDL-C individual data**

*p* values of MR-link after application to Lifelines individual level data and eQTLs from the BIOS cohort compared to the expected *p* value distribution (dashed line). (**A**) Using ordinary least squares (OLS) regression to solve equation 4 (**Methods**). (**B**) Using ridge regression to solve equation 5 (red) and ridge regression combined with *p* value calibration (green) (**Methods**) (**Supplementary Note**).

### Supplemental tables

**Supplemental Table 1**. **False positive rates and power of different MR methods and IV selection when no pleiotropy is simulated**.

Each row describes a simulation scenario and the corresponding detection rate using different approach for IV selection and different MR methods in a non-pleiotropic scenario ($b_U = 0.0$) (**Methods**). Simulation scenarios were varied by the (1) number of simulated causal SNPs (n_causal), (2) causal effect of the known exposure (exposure_1_causal) and (3) MR estimation method (estimation_method). The approach for IV selection (selection_method) varied between GCTA COJO (COJO) and p value clumping (clumped). Results are the median number of IVs identified by the selection method (median_ivs_identified), 'number_of_simulations' describes how often (out of 1,500 simulations) an estimation was made and finally how often a method identifies a significant effect at alpha = 0.05 (detection_rate). Note that detection rate is equivalent to false positive rate when exposure_1_causal = 0 and equivalent to detection power otherwise. See the **Supplementary Note** for an explanation of the MR methods MR-link OLS, MR-link ridge and MR-link ridge *p* calibrated.

*Excel file attached.*

**Supplemental Table 2**. **False positive rate and power of different MR methods when pleiotropy through linkage disequilibrium is simulated**.

Each row describes a simulation scenario and the corresponding detection rate for different MR methods in a pleiotropic scenario ($b_U = 0.4$) (**Methods**). Simulation scenarios were varied by the (1) the number of simulated causal SNPs (n_causal), (2) causal effect of the known exposure (exposure_1_causal) and (3) MR estimation method (estimation_method). Other columns indicate: the median number of IVs identified by GCTA-COJO (median_ivs_identified), how often (out of 1,500 simulations) an estimate was made (number_of_simulations) and how often a method identifies a significant effect at alpha = 0.05 (detection_rate). Note that detection rate is equivalent to false positive rate when exposure_1_causal=0 and equivalent to detection power otherwise. See the **Supplementary Note** for an explanation of the MR methods MR-link OLS, MR-link ridge and MR-link ridge *p* calibrated.

*Excel file attached.*

**Supplemental Table 3. False positive rate and power of different MR methods when pleiotropy through overlap is simulated**.

Each row describes a simulation scenario and the corresponding detection rate for different MR methods in a pleiotropic scenario ($b_U = 0.4$) and 10 simulated causal SNPs (**Methods**). The first columns indicate (1) the causal effect of the known exposure (exposure_1_causal), (2) the MR estimation method (estimation_method) and (3) the number of causal variants that overlap between the known- and pleiotropic exposure (overlapping_causal). The other columns indicate: the median number of IVs identified by the GCTA-COJO (median_ivs_identified), how often (out of 1,500 simulations) an estimate was made (number_of_simulations) and how often a method identifies a significant effect at alpha = 0.05 (detection_rate). Note that detection rate is equivalent to false positive rate when exposure_1_causal = 0 and equivalent to detection power otherwise. See the **Supplementary Note** for an explanation of the MR methods MR-link OLS, MR-link ridge and MR-link ridge *p* calibrated.

*Excel file attached.*

**Supplemental Table 4. Full results of the genes that were significant using MR-link and the eQTL from the BIOS cohort.**

In this table we list the full results for the 15 genes that pass significance in the MR-link analysis that used eQTLs from BIOS and the Lifelines cohort. We report the following information per gene: the Ensembl ID (ensg_id), gene ID (gene_id), causal estimate (estimated beta), standard error of the causal estimate (SE), *p* value of the estimate (calibrated_p), permuted *p* value (permuted_p) and finally the number of IVs identified by GCTA-COJO (IVs_identified). The causal estimate and the standard error refer to the uncalibrated pvalue.

| ensg_id | gene_id | Estimated beta | SE | calibrated_p | permuted_p | IVs_identified |
|---|---|---|---|---|---|---|
| ENSG00000245954 | *RP11-18H21.1* | 0.030 | 0.014 | 2.48E-07 | 0.004 | 2 |
| ENSG00000261087 | *KB-1460A1.5* | 0.154 | 0.062 | 5.55E-09 | 0.005 | 1 |
| ENSG00000068615 | *REEP1* | -0.022 | 0.010 | 5.72E-08 | 0.003 | 1 |
| ENSG00000002726 | *AOC1* | -0.009 | 0.004 | 1.18E-07 | 0.01 | 1 |
| ENSG00000222037 | *IGLC6* | -0.077 | 0.034 | 1.04E-07 | 0 | 3 |
| ENSG00000184292 | *TACSTD2* | -0.019 | 0.009 | 4.45E-07 | 0.003 | 2 |
| ENSG00000266852 | *MIR4482-1* | -0.107 | 0.044 | 7.41E-09 | 0 | 3 |
| ENSG00000138738 | *PRDM5* | 0.067 | 0.035 | 2.71E-06 | 0.001 | 4 |
| ENSG00000153157 | *SYCP2L* | -0.019 | 0.009 | 2E-07 | 0.001 | 2 |
| ENSG00000107731 | *UNC5B* | -0.012 | 0.005 | 2.13E-08 | 0.003 | 1 |
| ENSG00000211637 | *IGLV4-69* | 0.088 | 0.040 | 1.49E-07 | 0 | 1 |
| ENSG00000002933 | *TMEM176A* | -0.022 | 0.010 | 2.36E-07 | 0.004 | 3 |
| ENSG00000254030 | *IGLC5* | -0.053 | 0.020 | 7.75E-10 | 0 | 1 |
| ENSG00000165507 | *C10orf10* | -0.069 | 0.032 | 2.33E-07 | 0.001 | 1 |
| ENSG00000106565 | *TMEM176B* | -0.029 | 0.013 | 5.11E-08 | 0 | 4 |

**Supplemental Table 5. Significant genes identified in the MR-link analysis that used GTEx eQTL data.**

This table lists the full results for all genes found to be significant in the MR-link analysis that used GTEx eQTL summary statistics. It contains the following information per gene: the eQTL tissue (tissue), Ensembl ID (ensg_id), gene ID (gene_id), causal estimate (estimated_beta), standard error of the causal estimate (SE) , *p* value of the estimate (calibrated_p), permuted *p* value (permuted_p) and finally the number of IVs identified by GCTA-COJO (IVs_identified). The causal estimate and the standard error refer to the uncalibrated pvalue.

| tissue | ensg_id | gene_id | estimated_beta | SE | calibrated_p | permuted_p | IVs_identified |
|---|---|---|---|---|---|---|---|
| Liver | ENSG00000130202.5 | *PVRL2* | 0.318 | 0.052 | 3.24E-14 | 0 | 1 |
| Liver | ENSG00000134222.12 | *PSRC1* | -0.085 | 0.018 | 4.17E-09 | 0 | 1 |
| Liver | ENSG00000143126.7 | *CELSR2* | -0.099 | 0.024 | 7.04E-08 | 0 | 1 |
| Liver | ENSG00000134243.7 | *SORT1* | -0.087 | 0.019 | 6.15E-09 | 0 | 1 |
| Whole Blood | ENSG00000204920.6 | *ZNF155* | -0.069 | 0.034 | 9.8E-06 | 0.004 | 2 |
| Whole Blood | ENSG00000134222.12 | *PSRC1* | -0.157 | 0.069 | 1.69E-06 | 0 | 1 |

**Supplemental Table 6. Results of all other MR methods tested for the 15 genes identified using MR-link and the BIOS cohort**

This table lists the results of inverse variance weighting (IVW), MR-Egger, LDA-MR-Egger and MR-PRESSO for the 15 genes that pass significance in the MR-link analysis that used BIOS eQTL cohorts. Each row contains the following information: the Ensembl ID (ensg_id), gene ID (gene_id), method for causal estimation (method), causal estimate (estimated beta), standard error of the causal estimate (SE), *p* value of the estimate (p_val_estimated), and finally the number of IVs identified by GCTA-COJO (IVs_identified). Please note that in all cases, MR-PRESSO identified too many outliers and an estimate was not made.

| ensg_id | gene_id | method | estimated_beta | SE | p_val_estimated | IVs_identified |
|---|---|---|---|---|---|---|
| ENSG00000245954 | *RP11-18H21.1* | IVW | 0.016 | 0.005 | 0.000729 | 2 |
| ENSG00000261087 | *KB-1460A1.5* | IVW | 0.228 | 0.072 | 0.001402 | 1 |
| ENSG00000068615 | *REEP1* | IVW | -0.024 | 0.009 | 0.007291 | 1 |
| ENSG00000002726 | *AOC1* | IVW | -0.010 | 0.004 | 0.007236 | 1 |
| ENSG00000222037 | *IGLC6* | IVW | -0.042 | 0.011 | 9.09E-05 | 3 |
| ENSG00000222037 | *IGLC6* | MR-Egger | -0.086 | 0.014 | 0.103933 | 3 |
| ENSG00000222037 | *IGLC6* | MR-PRESSO | | | | 3 |
| ENSG00000222037 | *IGLC6* | LDA-MR-Egger | -0.079 | 0.027 | 0.212663 | 3 |
| ENSG00000184292 | *TACSTD2* | IVW | -0.011 | 0.003 | 8.15E-05 | 2 |
| ENSG00000266852 | *MIR4482-1* | IVW | -0.012 | 0.007 | 0.082145 | 3 |
| ENSG00000266852 | *MIR4482-1* | MR-Egger | -0.029 | 0.019 | 0.371586 | 3 |
| ENSG00000266852 | *MIR4482-1* | MR-PRESSO | | | | 3 |
| ENSG00000266852 | *MIR4482-1* | LDA-MR-Egger | -0.014 | 0.001 | 0.054 | 3 |
| ENSG00000138738 | *PRDM5* | IVW | 0.019 | 0.008 | 0.018 | 4 |
| ENSG00000138738 | *PRDM5* | MR-Egger | 0.015 | 0.007 | 0.171 | 4 |
| ENSG00000138738 | *PRDM5* | MR-PRESSO | | | | 4 |
| ENSG00000138738 | *PRDM5* | LDA-MR-Egger | 0.013 | 0.031 | 0.708 | 4 |
| ENSG00000153157 | *SYCP2L* | IVW | -0.013 | 0.003 | 7.59E-05 | 2 |
| ENSG00000107731 | *UNC5B* | IVW | -0.016 | 0.005 | 0.001 | 1 |
| ENSG00000211637 | *IGLV4-69* | IVW | 0.087 | 0.035 | 0.014 | 1 |
| ENSG00000002933 | *TMEM176A* | IVW | -0.009 | 0.004 | 0.045 | 3 |
| ENSG00000002933 | *TMEM176A* | MR-Egger | -0.001 | 1.85E-05 | 0.022 | 3 |
| ENSG00000002933 | *TMEM176A* | MR-PRESSO | | | | 3 |
| ENSG00000002933 | *TMEM176A* | LDA-MR-Egger | -0.004 | 0.001 | 0.159 | 3 |
| ENSG00000254030 | *IGLC5* | IVW | -0.040 | 0.016 | 0.010 | 1 |
| ENSG00000165507 | *C10orf10* | IVW | -0.071 | 0.030 | 0.018 | 1 |

| | | | | | | |
|---|---|---|---|---|---|---|
| ENSG00000106565 | *TMEM176B* | IVW | -0.010 | 0.004 | 0.010 | 4 |
| ENSG00000106565 | *TMEM176B* | MR-Egger | -0.014 | 0.008 | 0.201 | 4 |
| ENSG00000106565 | *TMEM176B* | MR-PRESSO | | | | 4 |
| ENSG00000106565 | *TMEM176B* | LDA-MR-Egger | -0.005 | 0.003 | 0.258 | 4 |
